# Attenuative effect of *Grifola frondosa* (maitake mushroom) on severe DSS-induced colitis in vitamin D-deficient mice

**DOI:** 10.1101/2025.04.04.647175

**Authors:** Miyu Nishikawa, Risa Miyagi, Yuki Kugimiya, Seita Chudan, Yukihiro Furusawa, Shinichi Ikushiro

## Abstract

**Scope:** Vitamin D deficiency, which has been a global health issue for decades, is involved in gut health and diseases. We examined the health benefits of *Grifola frondosa* (maitake mushroom), a potent dietary source of fungal vitamin D (D_2_), on DSS-induced colitis in vitamin D-deficient mice.

**Methods and results:** C57BL/6J mice were fed a control diet, vitamin D_3_-deficient diet (DD), maitake-fortified DD, or vitamin D_2_-fortified DD for 6 weeks. LC-MS/MS analysis demonstrated that maitake-fed mice showed an increased 25(OH)D_2_ alternative to 25(OH)D_3_ in plasma, as well as the mice fed an equivalent dose of vitamin D_2_. The mRNA expression profiles of vitamin D-responsive genes, including renal *Cyp24a1* and *Cyp27b1*, were normalized in the maitake-fed mice. Severe DSS-induced colitis observed in DD mice was attenuated in maitake-fed mice; the accumulation of immune cells in the colonic mucosa and protein expression of colonic claudin-2, a target gene of the vitamin D receptor, were comparable to that in control mice.

**Conclusion:** Dietary intake of maitake was effective in improving vitamin D status and biological function, demonstrating a potential attenuative effect on severe DSS-induced colitis in vitamin D_3_ deficient mice, as well as equivalent doses of vitamin D_2_.

## Introduction

Vitamin D exerts various biological effects, including calcium homeostasis, bone metabolism, anticancer, and immunomodulatory effects [1]. It exists in two forms, D_2_ and D_3_, which are the major forms in mushrooms and animals and differ in their side chains, with an additional CH2-group at C-24 and a double bond between C-22 and 23 for D_2_ (Figure 1A) [2, 3]. In humans, vitamin D_3_ is produced in the epidermal layer of the skin from 7-dehydrochoresterol by ultraviolet radiation. Dietary vitamin D, including D_2_ and D_3_, is bioavailable and maintains vitamin D levels in the body. 25-hydroxyvitamin D (25(OH)D), which is converted from vitamin D by CYP2R1 and CYP27A1 in the liver, is a major and long-life circulating metabolite of vitamin D and a reliable biomarker showing sufficient vitamin D status in the body [4, 5]. According to the guidelines of the American Endocrine Society, vitamin D sufficient status is defined as follows depending on serum 25(OH)D levels: < 20 ng/mL; vitamin D deficiency, 21 to 29 ng/mL; vitamin D insufficiency [6]. To maintain adequate levels (40–60 ng/mL) of 25(OH)D, a daily intake of 400–800 IU of vitamin D is recommended, depending on age and sex [7].

**Figure 1.**
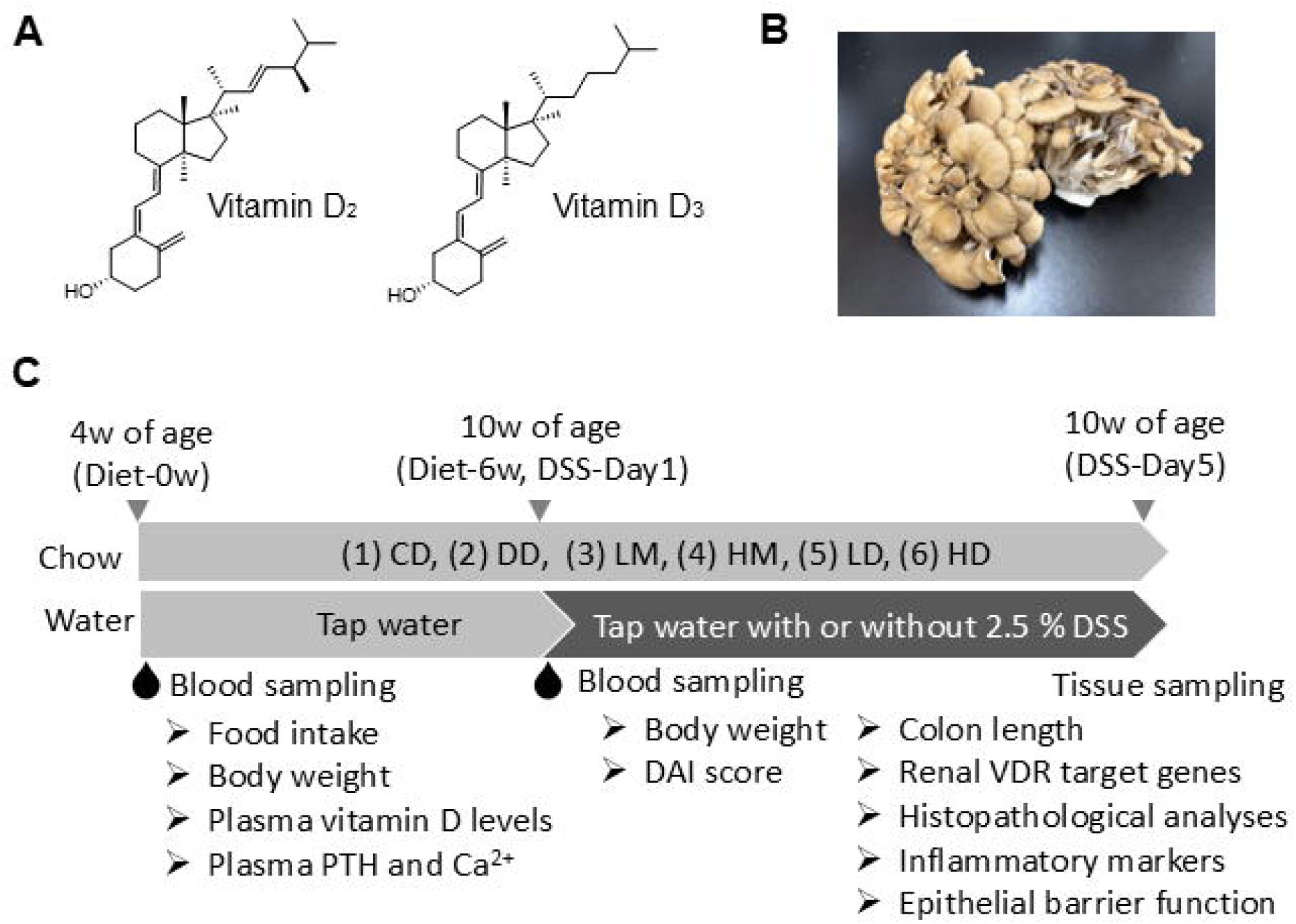
Overview of the present study. (A) Chemical structure of vitamin D_2_ and D_3_. (B) Picture of the maitake used in this study. (C) Study design of the DSS-induced colitis model with different vitamin D sufficient status.

Despite the endogenous and dietary availability of vitamin D, vitamin D insufficiency and deficiency have been a worldwide health issue for decades [8], posing a potential risk for diverse types of related diseases, such as osteoporosis, cancer, and other chronic and metabolic diseases such as obesity [9]. Screening for vitamin D deficiency in U.S. children reported that 9% of the pediatric population had vitamin D deficiency and 61% had vitamin D insufficiency [10]. Although vitamin D-fortified foods, such as milk and cereals, are commonly consumed in the U. S., only 4% of the subjects had consumed 400 IU of vitamin D per day for the past 30 days [10]. Vitamin D deficiency appears to be more severe problem in Japan. In addition to excessive avoidance of sunlight exposure, lifestyle changes such as spending less time outdoors resulting from the COVID-19 pandemic are also probable causes of severe vitamin D deficiency. The prevalence of vitamin D deficiency in Japanese young women (21–25 years) increased to 90.0 %, compared to 38.5% in 2016 [11]. The mass screening conducted in Tokyo from 2019 to 2020 demonstrated a high prevalence of vitamin D insufficiency in healthy subjects: 98% of the subjects showed vitamin D insufficiency or deficiency [12]. The “vitamin D deficiency pandemic” may accordingly increase health hazards [8]; thus, novel nutritional strategies to improve vitamin D status are needed for human health promotion.

Unfortunately, few natural foods contain vitamin D, although some types of edible mushrooms or fatty fish such as trout and salmon are useful sources of vitamin D_2_ and D_3,_ respectively. In the U. S., milk is fortified with vitamin D, being the major vitamin D source [13, 14]. However, vitamin D fortification of food products is not prevalent in Japan. Thus, natural food ingredients such as salmon and edible mushrooms are potent vitamin D sources. *Grifola frondose* (Figure 1B), known as maitake in Japanese, contains higher amounts of vitamin D_2_ than other mushrooms [15]. Although dried shiitake mushroom (12.7 μg vitamin D_2_ /100 g dried weight) has been a popular vitamin D_2_ source in the past, improved food transportation and preference for time-saving cooking in recent years increased consumption of raw shiitake mushroom, which contains less vitamin D_2_ than dried one. Today, maitake mushrooms have become popular and easily available in grocery stores in Japan as agricultural cultivation has developed since the 1980s. Despite the nutritional potency of maitake mushrooms as a novel vitamin D source in Japan, few studies have demonstrated the bioavailability and biological functions of the vitamin D_2_ contained in maitake. In fact, screening for vitamin D sufficiency in more than 5,000 Japanese subjects demonstrated that most had undetectable concentrations of 25(OH)D_2_ [12]. While it has been considered that vitamin D_2_ content in maitake mushroom is 4.9 μg per 100 g fresh weight, higher vitamin D_2_-contained maitake, which contains more than 20.0 μg vitamin D_2_ in 100 g, is also commercially available in recent years. Therefore, we focused on the health benefits of maitake through the vitamin D_2_ action.

Vitamin D is sequentially converted to its final active form 1α,25-dihydroxivitamin D (1,25(OH)_2_D), which exerts transcriptional activity on target genes via its high binding affinity to the vitamin D receptor (VDR). Thus, a lack of vitamin D signals, including vitamin D deficiency and dysfunctional mutations in the VDR, causes undesirable health outcomes such as bone disorders. Furthermore, vitamin D signaling has been implicated in gastrointestinal health. Global or tissue-specific deletion of VDR in intestinal or colorectal epithelial cells resulted in severe progression of dextran sodium sulfate (DSS)-induced colitis in mice [16–20]. Analytical epidemiological studies in recent years have suggested that vitamin D deficiency or insufficiency is associated with disease activity in patients with inflammatory bowel diseases (IBD), including ulcerative colitis and Crohn’s disease [21–26], although vitamin D deficiency may result from malabsorption of dietary vitamin D due to the involvement of the upper gastrointestinal tract in Crohn’s disease patients [21]. Although the incidence and prevalence of IBD have increased globally in the 21st century, very few medicinal approaches are currently effective. In the present study, we aimed to investigate the effects of maitake intake on vitamin D sufficiency and intestinal health. We examined the effects of maitake feeding on (1) alternative improvement of vitamin D status in vitamin D_3_ -deficient mice, and (2) the progression of DSS-induced colitis.

## Materials and Methods

### 1. Materials

Commercial-grade maitake was kindly provided by HOKUTO Corporation (Nagano, Japan). Dextran sodium sulfate (DSS, MW36,000-50,000) was obtained from MB Biomedical (Irvine, CA, U.S.). D6-25-hydroxyvitamin D_3_ (26,26,26,27,27,27-D6, d6-25(OH)D_3_) and LC-MS-grade organic solvents were purchased from Sigma-Aldrich (St. Louis, MO, U.S.). 4-[2-(6,7-dimethoxy-4-methyl-3-oxo-3,4-dihydroquinoxalyl)ethyl]-1,2,4-triazoline-3,5-dione (DMEQ-TAD) and authentic standards of 25(OH)D_2_ and 25(OH)D_3_ were purchased from FUJIFILM Wako Pure Chemical (Osaka, Japan). All other chemicals were commercially available and of the highest quality.

### 2. Measurement of vitamin D_2_ concentration in maitake mushrooms

The vitamin D_2_ concentration in maitake was determined using HPLC at Japan Food Research Laboratories (Tokyo, Japan). The vitamin D_2_ concentration of pooled maitake was 23.7 μg/100 g fresh weight and vitamin D_3_ was under-detectable (LOQ: 0.7 μg/100 g).

### 3. Animals and diet

Control diet (D10012GM, AIN-93G based) and vitamin D_3_-deficient diet (D10073001M, AIN-93G based) were purchased from Research Diets (New Brunswick, NJ, U.S.). Male C57BL/6J mice (3 weeks old) were supplied from Japan SLC (Hamamatsu, Japan) and housed in a controlled room temperature (22 to 26 °C), and in 50 to 55% humidity with a 12 h light/dark cycle. They were given a control diet and water ad libitum for one week for acclimatization.

In this study, we prepared six types of diets as follows: control diet (CD), vitamin D_3_-deficient diet (DD), DD containing low dose (1,000 IU eq. vitamin D_2_/kg) of maitake powder (LM), DD containing high dose (2,000 IU eq. vitamin D_2_/kg) of maitake powder (HM), DD containing a dose of pure vitamin D_2_ equivalent to that in LM (LD), and DD containing a dose of pure vitamin D_2_ equivalent to that in HM (HD). Maitake was freeze-dried until its weight was reduced to one-tenth of its fresh weight and then ground into a powder. Maitake powder was added to the vitamin D_3_-deficient diet and mixed thoroughly to prepare a maitake-containing diet. Finally, LM and HM contained 1,000 and 2,000 IU vitamin D_2_/kg diet, respectively. Doses of pure vitamin D_2_ (FUJIFILM Wako Pure Chemical) equivalent to LM and HM were also added to the vitamin D_3_-deficient diet and mixed well, which formed LD and HD, respectively. The diet composition is summarized in Table S1.

### 4. Experimental design

At four weeks of age, conditioned mice were randomly assigned to six diet groups: CD, DD, LM, HM, LD, and HD (each diet group was further divided into DSS- and + groups, Figure 1C). After raising the mice on the test diet for 6 weeks, blood was collected from the jugular vein prior to the DSS challenge, as the blood component after the challenge was affected by ruminal bleeding. Blood was centrifuged at 3,000 × *g* for 10 min, and the resulting plasma sample was stored at −80 °C for subsequent experiments.

The DSS+ group was given drinking water containing 2.5 % DSS for 4 days, with definition of first day as Day1. On the fourth day (Day5) of DSS challenge, the mice were sacrificed by incision of the inferior vena cava under isoflurane anesthesia to remove the blood. Tissues, including the kidney and proximal colorectal mucosa, were collected and frozen in liquid nitrogen. The distal colon was incised, rinsed with cold PBS twice, and fixed with 4% paraformaldehyde. The disease activity index (DAI) was assessed based on body weight gain, bleeding, and colon length according to a previous study [31]. All the biological samples were stored at −80 °C for subsequent experiments.

All experimental protocols involving animals were approved by the Animal Research and Ethics Committee of Toyama Prefectural University (approval number: R5-13).

### 5. Dosage information

Vitamin D_2_ concentration in the test diet was as follows: 1,000 IU (25 μg) vitamin D_2_/kg diet in LM and LD group, 2,000 IU (50 μg) vitamin D_2_/kg diet in HM and HD group. We chose these doses to make vitamin D concentration in rescue diet equivalent to control diet (1,000 IU vitamin D_3_/kg), with consideration for bioavailability of maitake-derived vitamin D. As food intake of mice during the test period was approximately 3 g (Figure S3), they are estimated to take 3 IU (7.5 ng) or 6 IU (150 ng) vitamin D_2_/day. Considering the body weight of mice was around 15 to 20 g, the human equivalent dose (HED) of vitamin D_2_ is 9,000 or 18,000 IU/day, in the case of 60 kg body weight. Although it exceeds the recommended daily amount of vitamin D, 400–800 IU [7], the primary objective of this study is to examine the compensatory vitamin D action of maitake in DD mice.

### 6. LC-MS/MS analysis of plasma 25(OH)D_2_ and 25(OH)D_3_

Plasma concentrations of 25(OH)D_2_ and 25(OH)D_3_ were measured using LC-MS/MS as described previously, with slight modifications [27, 28]. Quantitative analysis was carried out using MS/MS-multiple reaction monitoring (MRM) of the precursor/product ion for DMEQ-TAD-25(OH)D_3_ (*m*/*z*: 746.5/468.1), DMEQ-TAD-25(OH)D_2_ (*m*/*z*: 758.6/468.1), and DMEQ-TAD-*d*_6_-25(OH)D_3_ (*m*/*z*: 752.5/468.1) with a dwell time of 200 ms (see Figure S1 and Table S2).

### 7. Measurement of vitamin D-responsive parameters in plasma

Plasma concentrations of 1,25(OH)_2_D_2_ and 1,25(OH)_2_D_3_ were determined as total 1,25(OH)_2_D by using the 1, 25-(OH)_2_ Vitamin D ELISA Kit (Immundiagnostik, Bensheim, Germany) as described previously [27, 28]. Plasma parathyroid hormone (PTH) levels were determined using a mouse PTH 1-84 ELISA kit (Immutopics, San Clemente, CA, U.S.), following the manufacturer’s protocol. Plasma ionized calcium (Ca^2+^) concentrations were determined using the Calcium Test E Wako (Wako, Tokyo, Japan).

### 8. Real-time quantitative PCR

Total RNA was isolated from the kidney using ISOGEN II (Nippon Gene, Tokyo, Japan). cDNA was synthesized using the PrimeScript RT Master Mix (Perfect Real-Time) (TaKaRa, Otsu, Japan). Real-time PCR was carried out with an Applied Biosystems 7500 Real-Time PCR System using TB Green Premix Ex Taq II (TaKaRa) for the reaction reagent. The mRNA expression of *Cyp24a1* and *Cyp27b1* was determined with the ΔΔCt method using β*-actin* as an internal control (Table S3).

### 9. Histological analysis

Distal colonic sections were prepared from the paraffin-embedded samples at the Cell Technology Laboratory (Sapporo, Japan). The deparaffinized sections were stained with hematoxylin and eosin. For immunofluorescence analysis, deparaffinized colonic sections were microwaved for 15 min in 10 mM sodium citrate buffer. After blocking with UltraCruz Blocking Reagent (Santa Cruz Biotechnology, Dallas, TX, U.S.), the sections were incubated with an anti-S100A9 antibody (Proteintech, Rosemont, IL, U.S.). After the sections were washed with PBS containing 0.05% Tween-20 for 5 min thrice, they were incubated with FITC-conjugated anti-rabbit antibody for 2 h in the dark. Similarly, the sections were washed three times, mounted with VECTASHIELD Mounting Medium with DAPI (Vector Laboratories, Inc. (Burlingame, CA, U.S.), and then analyzed using an OLYMPUS IXplore Fluorescence Microscope (OLYMPUS, Tokyo, Japan). The S100A9 positive cells were measured using Image J software (version 1.54).

### 10. Immunoblotting

The proximal colonic mucosa was homogenized and sonicated in cold RIPA buffer containing 1% proteinase inhibitor cocktail for use with mammalian cells and tissues (Nacalai Tesque, Kyoto, Japan). After centrifugation at 8,000 × *g* for 10 min, the protein concentration in the supernatant was measured using a BCA protein assay kit (Nacalai Tesque). The protein sample containing Sample Buffer Solution with Reducing Reagent (6×) for SDS-PAGE (Nacalai Tesque) was incubated at 95 ℃ for 5 min.

The SDS-PAGE was performed using p-PAGEL (ATTO, Tokyo, Japan) in EzRun T (ATTO) electrophoresis running buffer. After the separated proteins were blotted onto a nitrocellulose membrane, the membrane was blocked with Blocking One (Nacalai Tesque) for 15 min and incubated with an anti-claudin-2 antibody (Santa Cruz Biotechnology) overnight. After washing with PBS containing 0.1% tween 20 thrice, the membrane was incubated with anti-mouse IgG AP-linked antibody (Cell Signaling Technology, Danvers, MA, U.S.) for 2 h. The membrane was washed with TBS containing 0.1% Tween 20 thrice and then incubated with BCIP-NBP solution (Nacalai Tesque) for coloring reaction. The band area was measured using Image J software (version 1.54) and the expression of Cld-2 was calibrated using β-actin.

### 11. Statistical analysis

The analysis was conducted using the IBM SPSS Statistics software (version 25). One-way analysis of variance (ANOVA) was performed. Differences were considered significant at *p* < 0.05 using two-tailed Tukey’s t-test.

## Results

### 1. Effect of maitake feeding on circulating vitamin D status in vitamin D_3_-deficient mice

Both vitamins D_2_ and D_3_ are individually converted to 25(OH)D_2_ and D_3_ by CYP2R1 and CYP27A1 in the liver, respectively, which is a biomarker of sufficient vitamin D status. Thus, we measured plasma 25(OH)D_2_ and D_3_ levels in mice fed the following test chow: control diet (CD), vitamin D_3_-deficient diet (DD), low maitake diet (LM), high maitake diet (HM), low vitamin D_2_ diet (LD), and high vitamin D_2_ diet (HD). For 25(OH)D_3_, the plasma concentration significantly decreased in all dietary groups, except for the CD group, after 6 weeks of the test diet period (Figure 2A). The 25(OH)D_2_ concentration in CD and DD mice was 0.8 ± 0.37 and 0.9 ± 0.40 ng/mL, respectively (Figure 2A), suggesting that the effect of vitamin D_2_ in AIN-93G diet formula is almost negligible. In DD mice, 25(OH)D_3_ levels rapidly decreased during the first week of DD feeding (Figure S2), indicating that the vitamin D_3_-deficient mouse model used in this study maintained vitamin D_3_ deficiency for 5 weeks before the subsequent DSS challenge. In contrast to CD and DD mice, the 25(OH)D_2_ levels in LM, HM, LD, and HD mice were significantly elevated, reaching approximately 30 ng/mL, which is categorized as vitamin D sufficiency (Figure 2A). These results indicate that maitake-derived vitamin D_2_ is as bioavailable as purified vitamin D_2_. Thus, total plasma 25(OH)D in LM, HM, LD, and HD mice, which was calculated as the sum of the concentrations of 25(OH)D_2_ and 25(OH)D_3_, was significantly higher than that in DD mice (Figure 2A). No significant differences in food intake or body weight were observed during the test diet period (Figure S3).

**Figure 2.**
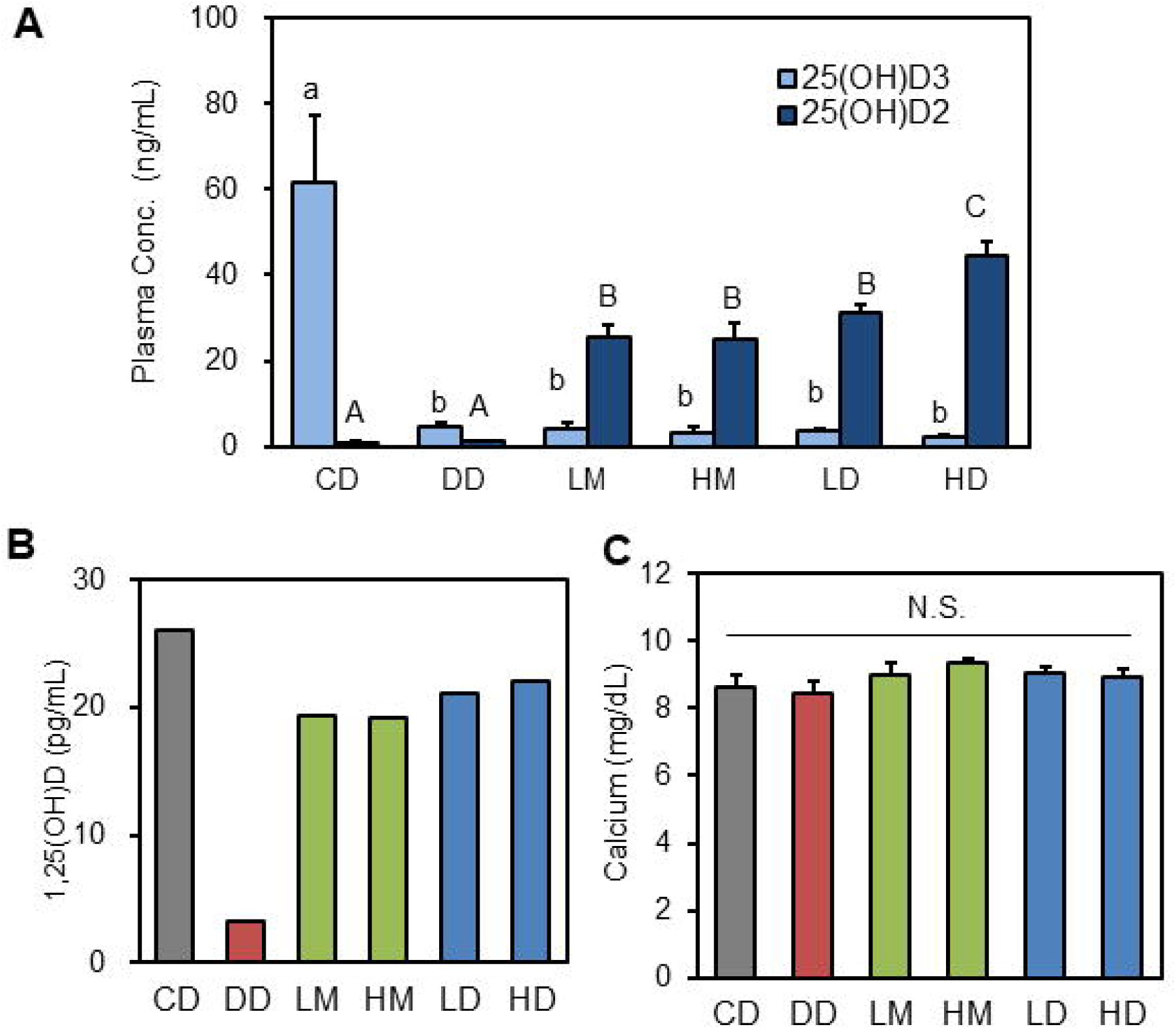
Plasma vitamin D status in mice fed a vitamin D-controlled diet for 6 weeks. (A) Plasma concentration of 25(OH)D_2_ and 25(OH)D_3_, (B) plasma 1,25(OH)_2_D concentration, and (C) plasma Ca^2+^ level in six diet groups after 6 weeks. The values are represented as mean ± SEM (n = 6–8 mice/group) in A and C, while the values are represented as the mean of technical duplicate using pooled sample (n = 6 mice/group) in B. One-way ANOVA followed by post-hoc Tukey’s t-test was performed in A and C. Without common letters (A vs B vs C, and a vs b): *p* < 0.05. N.S.: no significant difference.

The concentration of plasma 1,25(OH)_2_D, the biologically active form of vitamin D, was determined by ELISA using an antibody that reacts with both 1,25(OH)_2_D_2_ and D_3_. As expected, the plasma levels of 1,25(OH)_2_D (total concentrations of 1,25(OH)_2_D_2_ and D_3_) were significantly lower in DD mice, whereas the 1,25(OH)_2_D levels in LM, HM, LD, and HD mice recovered to a level equivalent to that in CD mice (Figure 2B). Severe vitamin D deficiency often results in hypocalcemia, as intestinal calcium absorption and renal calcium reabsorption are promoted by transient receptor potential vanilloids, such as TRPV5 and 6, which are well-known target genes of VDR. However, there was no significant difference in plasma calcium concentration among the test diet groups in the present study (Figure 2C).

These results strongly suggest that maitake can improve the circulating vitamin D status in vitamin D_3_-deficient mice, similar to pure vitamin D_2_. Maitake or purified vitamin D_2_-fortification also improved the level of the active form of vitamin D, 1,25(OH)_2_D, indicating that the major form of that is 1,25(OH)D_2_. Regarding 1,25(OH)_2_D quantification, the LC-MS approach was adequate for determining the individual concentrations of 1,25(OH)_2_D_2_ and D_3_. The reason for using ELISA with an antibody reacting to both 1,25(OH)_2_D_2_ and D_3_ is that we could not prepare sufficient plasma samples from mice for LC-MS, which requires a large sample volume owing to its lower sensitivity. However, the elevation of 25(OH)D_2_ instead of 25(OH)D_3_ in mice fed maitake or vitamin D_2_ strongly indicates that the recovered 1,25(OH)_2_D in them was mostly derived from 1,25(OH)_2_D_2_. Further detailed analyses are required to confirm the generation of 1,25(OH)_2_D_2_ as an alternative to 1,25(OH)_2_D_3_ in the maitake-fortified mice.

### 2. Effect of maitake feeding on the expression of vitamin D-responsive genes in vitamin D_3_-deficient mice

Decreased plasma 1,25(OH)_2_D concentration due to vitamin D_3_-deficient feeding was recovered in LM and HM mice as well as in LD and HD mice. We examined the mRNA expression profile of the vitamin D-responsive genes in the kidney, which is a major target organ of vitamin D. *Cyp24a1*, the expression of which is robustly induced by 1,25(OH)D-bound VDR, is involved in the negative feedback of vitamin D status by mediating C-24 or C-23 hydroxylation of 1,25(OH)_2_D and 25(OH)D (Figure 3A). The mRNA expression of renal *Cyp24a1* in DD mice was lower than that in CD mice, although this difference was not significant (Figure3B). Furthermore, the decreased *Cyp24a1* expression in DD mice was recovered in LM and HM mice in a dose-dependent manner up to a comparable level of CD and was further induced in LD and HD mice compared to CD mice.

**Figure 3.**
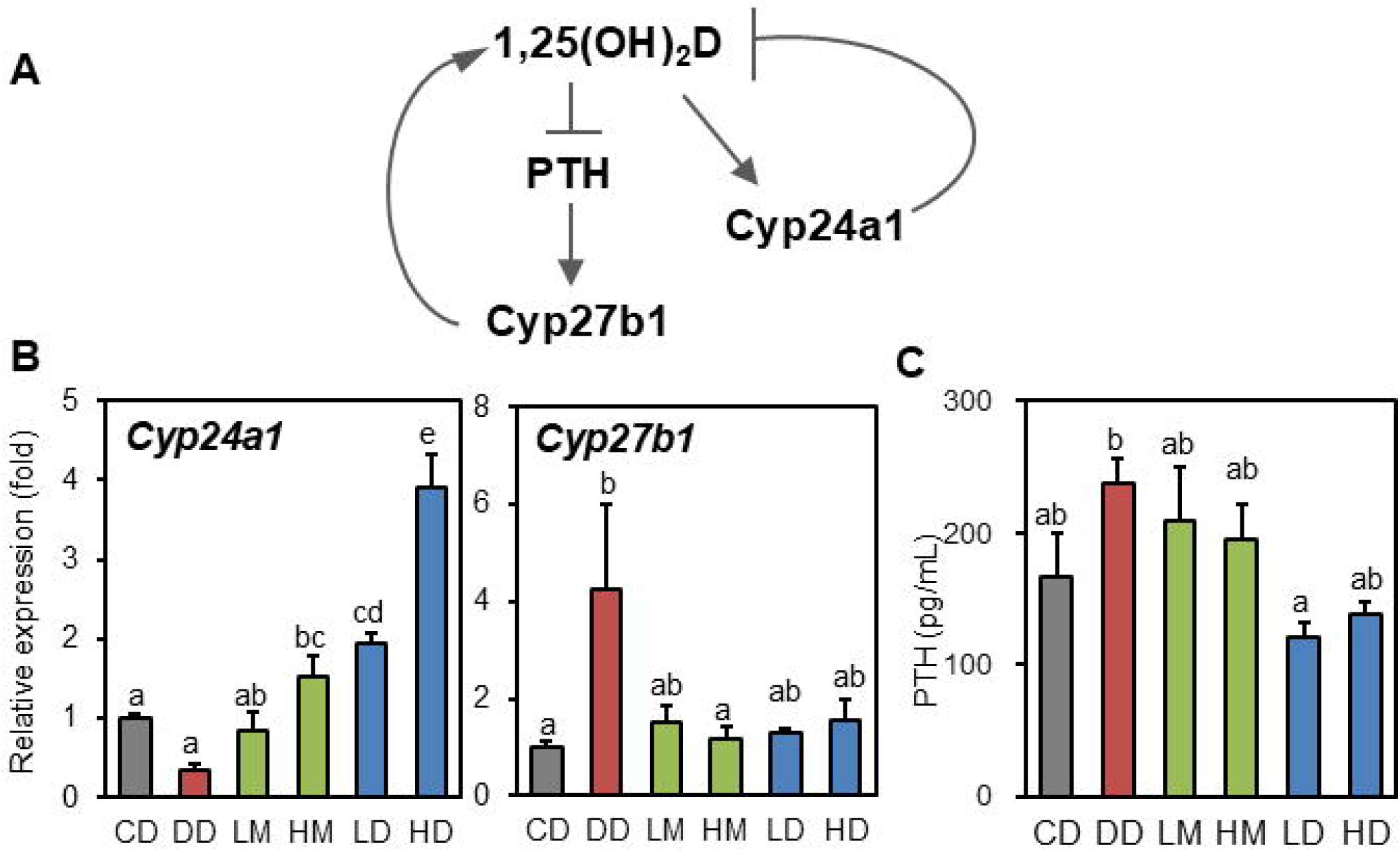
Biological responses to vitamin D after feeding with a vitamin D-controlled diet. (A) Biological response of *Cyp24a1*, *Cyp27b1*, and PTH according to the plasma concentration of 1,25(OH)_2_D. (B) Relative mRNA expression of *Cyp24a1* and *Cyp27b1* in the kidney, and (C) plasma PTH concentration in six diet groups of mice after 6 weeks. The values are represented as mean ± SEM (n = 5 and 4 mice/group in B and C, respectively). Without common letters: *p* < 0.05 by one-way ANOVA followed by post-hoc Tukey’s t-test.

The transcriptional response of renal *Cyp27b1* to 1,25(OH)_2_D was opposite to that of *Cyp24a1*, whose expression was induced under vitamin D-deficient conditions via parathyroid hormone (PTH) signaling (Figure 3A). Renal *Cyp27b1* expression in DD mice was significantly higher than that in CD mice, whereas its expression in LM, HM, LD, and HD mice was equivalent to that in CD mice (Figure 3B). The secretion of PTH, which is involved in vitamin D-linked calcium homeostasis, is triggered by decreased Ca^2+^ and 1,25(OH)_2_D levels in the blood in a direct or indirect manner (Figure 3A). The plasma PTH level of the DD mouse model increased, whereas maitake or pure vitamin D_2_ fortification normalized the PTH levels (Figure 3C). These biological responses were dependent on plasma 1,25(OH)D_2_ levels (Figure 2B), indicating that maitake-derived vitamin D_2_ is biologically functional.

### 3. Impact of maitake-feeding on the pathological index of DSS-induced colitis in vitamin D_3_-deficient mice

Decreased vitamin D signals, including dietary vitamin D deficiency and genetic VDR deletion, exacerbate the disease score in DSS-induced colitis [12, 17, 19, 29]. Hence, we examined the protective effects of maitake supplementation on the development of DSS-induced colitis in vitamin D_3_-deficient mice. Prior to the experiment, DSS treatment conditions for the colitis model were optimized so that the disease score in CD mice was mild because the DSS condition that showed a progressed disease score in CD mice often resulted in increased mortality of the DD mice during this period.

Under optimized conditions, DD mice showed a severe colitis index compared to CD mice after DSS challenge for 5 days, which is consistent with a previous report [29]. The body weight of DD mice significantly decreased on the 5th day of the DSS challenge, while that of CD mice did not change and even slightly increased (Figure 4A). The weight loss during the DSS challenge was partially recovered in LM mice compared to DD mice, and that in HM, LD, and HD mice was comparable to that in CD mice, which maintained weight gain during the DSS challenge (Figure 4A). Colonic shortening, which is associated with disease severity in DSS-induced colitis models [30], was more severe in DD mice than in CD mice, with severe bleeding in the colonic lumen (Figure 4B). In vitamin D_2_-rescued mice in the LM, HM, LD, and HD groups, the degree of colon shortening was comparable to that in CD mice (Figure 4B and 4C). The DAI score, which is a useful indicator of colitis progression [31], was higher in DD mice compared to CD mice, while the score in maitake or vitamin D_2_ fortified mice against CD mice was equivalent or even improved (Figure 4D). Although the intake of DSS solution per bodyweight was slightly higher in LM group (Figure S4), severity of the colitis was associated with vitamin D sufficient status.

**Figure 4.**
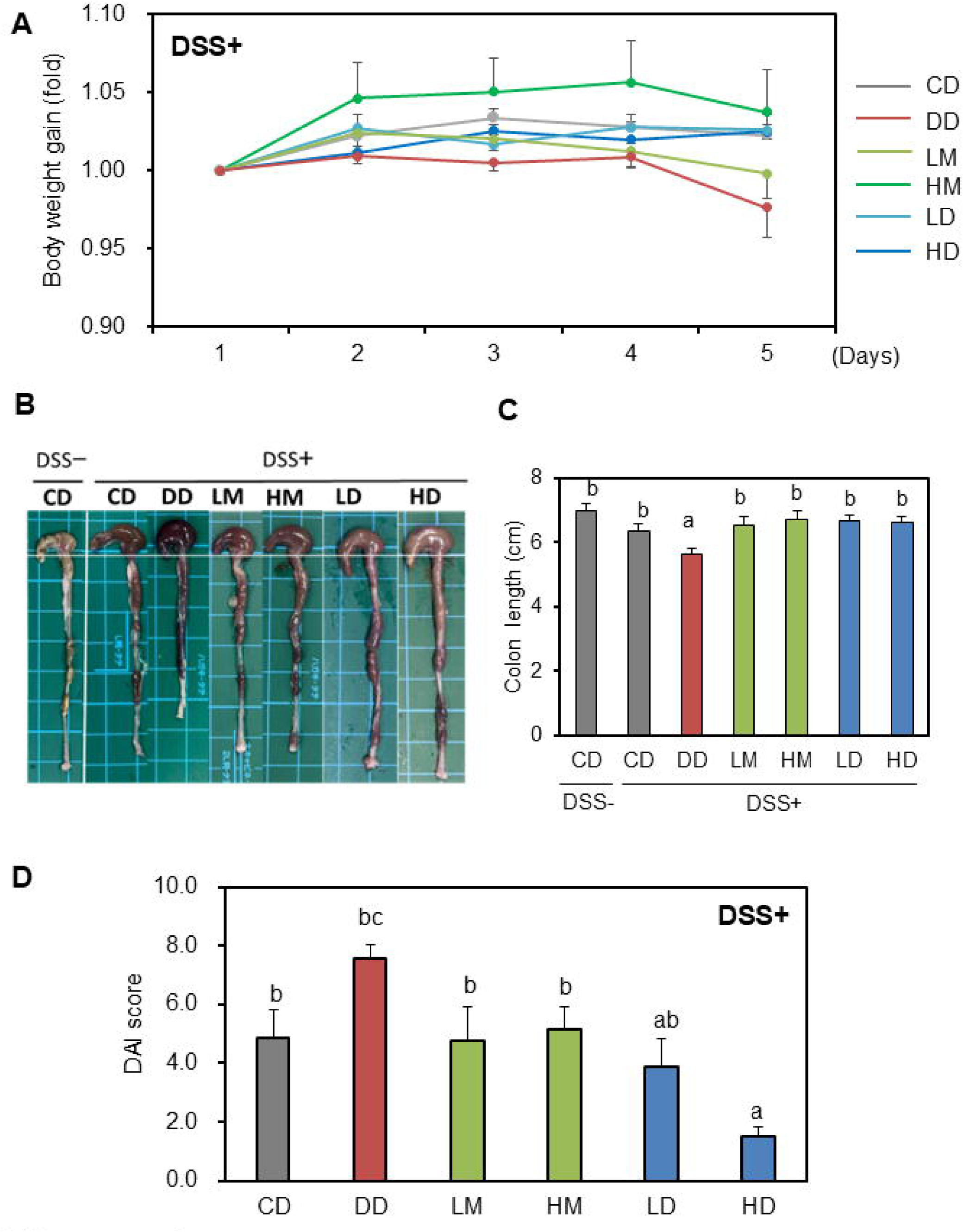
Disease activities of DSS-induced colitis mice fed a vitamin D-controlled diet. (A) Body weight gain during DSS-challenge. (B) Comparison of the colon appearance on day 5 of the DSS challenge. (C) Colon length of the colitis and non-colitis model raised with a vitamin D-controlled diet. *; *p* < 0.05 by one-way ANOVA followed by post-hoc Tukey’s t-test. (D) DAI score of the colitis models. The values are the means ± SEM (n = 6–8 mice/group in A, C, D). Without common letters: *p* < 0.05 by one-way ANOVA followed by post-hoc Tukey’s t-test.

Histological analysis also demonstrated a protective effect of maitake feeding against histological injury to the colonic mucosa. The H&E images of DD mice showed severe infiltration of immune cells into the mucosal layer, whereas vitamin D_2_-rescued mice, such as LM, HM, LD, and HD, showed drastically recovered epithelial structures with decreased lymphocyte infiltration (Figure 5A). Calprotectin, a heterotetramer composed of the S100A8 and A9 subunits and involved in wound healing, is a useful biomarker for IBD diagnosis [32–34]. In CD mice, S100A9-positive cells significantly increased in the colonic epithelium following DSS treatment (Figure 5A and 5B). In colitis mice, the number of positive cells increased further in DD mice, whereas it decreased in LM, HM, LD, and HD mice (Figure 5A and 5B). Thus, maitake feeding attenuated severe DSS-induced colitis in vitamin D_3_-deficient mice, similar to purified vitamin D_2_-feeding.

**Figure 5.**
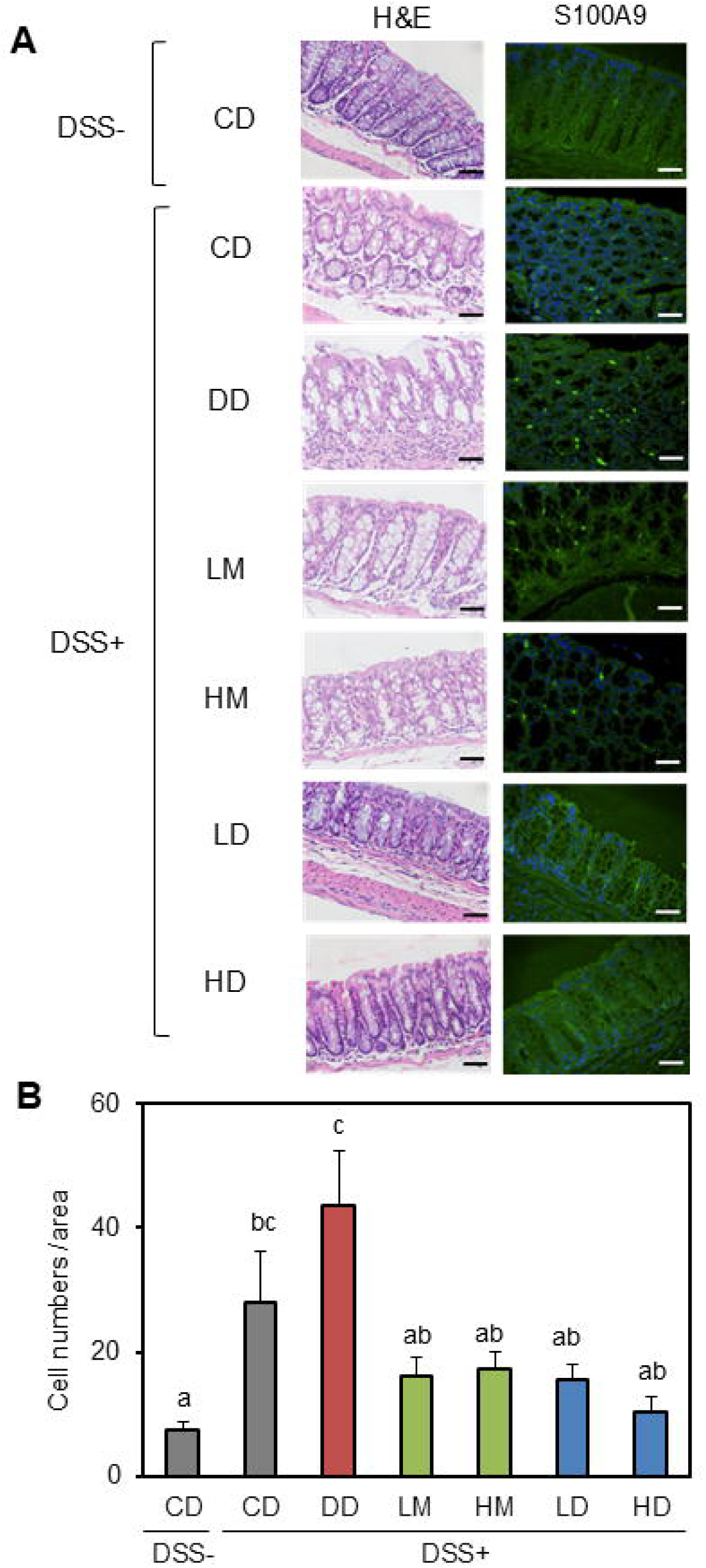
Histological evaluation of colonic injury in colitis or non-colitis models. (A) Representative image of colonic mucosa after H&E staining, and immuno-fluoresceine analysis of S100A9. Scale bar; 50 μm. (B) Quantitative analysis of S100A9-positive cells in the section. The cell numbers were calculated as the total amount of three independent areas in the same section. The values are then represented as the means ± SEM (n = 6–8 mice/group). Without common letters: *p* < 0.05 by one-way ANOVA followed by post-hoc Tukey’s t-test.

### 4. Impact of *G. frondosa* on the intestinal epithelial barrier in dietary vitamin D_3_-deficient mice

The expression of claudin-2 (Cld-2), which is a leaky tight junction protein and mediates paracellular cation transport [35, 36], is robustly upregulated in IBD patients [37, 38, 39, 40]. Cld-2 is also a direct target of VDR [41]. Vdr deletion in mice led to a robust upregulation of colonic Cld-2 under inflammatory conditions associated with more severe inflammation, whereas a lack of VDR downregulated Cld-2 expression under non-inflammatory conditions (Figure 6A) [20].

**Figure 6.**
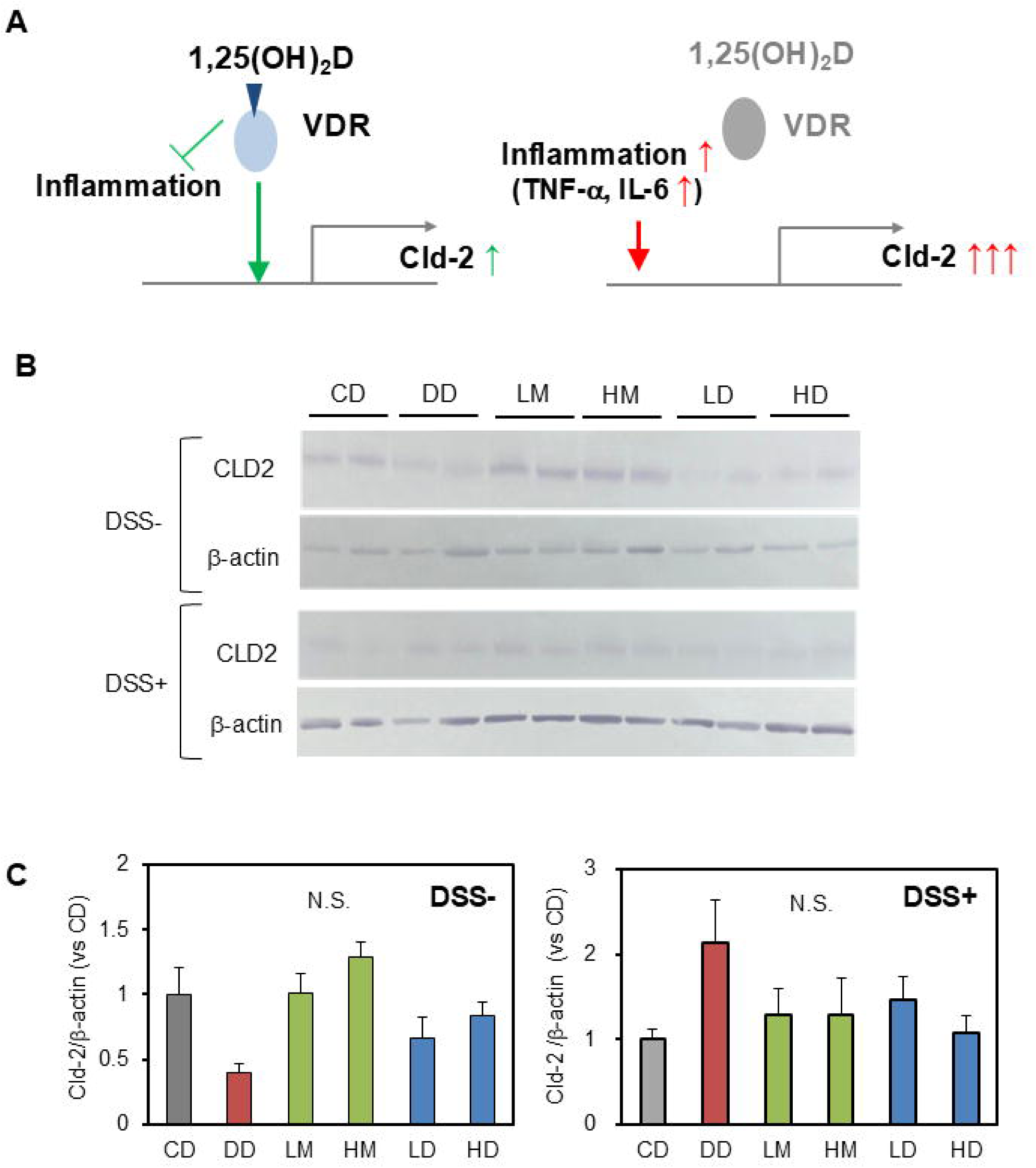
Effect of vitamin D status on colonic epithelial barrier. (A) Proposed regulation of Cld-2 in inflammatory intestinal epithelial cells through direct or indirect VDR-dependent signaling [20]. (B) Representative images of immunoblotting for Cld-2 and β-actin in the colonic mucosa (n=2 mice/group). (C) Relative protein expression in Cld-2. Values represent the means ± SEM (n = 6–8 mice/group). N.S.: no significant difference by one-way ANOVA followed by post-hoc Tukey’s t-test.

Thus, we examined whether the attenuative effect of maitake on DSS-induced colitis was mediated by regulation of Cld-2. As expected, Cld-2 expression in DD mice showed a decreasing tendency compared to CD mice under normal conditions, whereas it was comparable to CD mice in vitamin D_2_-rescued mice, such as LM, HM, and HD (Figure 6B and 6C). In contrast to normal conditions, Cld-2 expression in the colitis model tended to increase in DD mice (Figure 6B and 6C). Even under colitis conditions, the expression levels of Cld-2 in the LM, HM, LD, and CD mice were comparable to those in the CD mice (Figure 6B and 6C). The gut permeability of FITC-dextran also showed an increased tendency in DD mice compared to that in CD mice under colitis conditions (Figure S5). These results suggest that maitake feeding exerts a protective effect against severe DSS-induced colitis in vitamin D_3_-deficient mice through vitamin D signaling, which is involved in gut permeability via the regulation of Cld-2 expression. Although multiple physiological roles of vitamin D in gut health have been reported, including immunomodulatory and calcemic effects, gut permeability through Cld-2 may be one of the direct actions of maitake-derived vitamin D_2_ in colonic epithelial cells.

## Discussion

We demonstrated that maitake-fortified feeding was effective in improving vitamin D status in vitamin D_3_-deficient mice. The first notable finding of this study was that the bioaccessibility of vitamin D_2_ in maitake was comparable to the equivalent dose of purified vitamin D_2_, both of which increased 25(OH)D_2_ but not 25(OH)D_3_ in the plasma, associated with elevated 1,25(OH)_2_D (Figure 2A and 2B).

In addition to its good bioavailability, we demonstrated, for the first time, that maitake-derived vitamin D_2_ is biologically functional, which is assumed to be mediated by VDR-bound 1,25(OH)_2_D_2_. The mRNA expression profiles of vitamin D-responsive genes, including *Cyp24a1* and *Cyp27b1*, which were downregulated and upregulated in vitamin D_3_-deficiemt mice, respectively, were normalized to those in maitake-fed and purified vitamin D_2_-fed mice (Figure 3B). Finally, we demonstrated the potential attenuative effects of maitake on the severe disease activity of DSS-induced colitis in vitamin D_3_-deficient mice. Although vitamin D_3_-deficient mice in the DD group showed a more severe colitis index trend than vitamin D_3_-sufficient mice in the CD group, the vitamin D-rescued group using maitake or purified vitamin D_2_ showed an improved index compared to vitamin D_3_-deficient mice (Figure 4 and 5). The effect of vitamin D-rescue on the colitis index showed a similar tendency between maitake and equivalent purified vitamin D_2_-fed mice, strongly suggesting that these physiological activities are mediated by vitamin D signaling-derived dietary vitamin D_2_.

Although gut permeability in the colitis model with a vitamin D_3_-deficient diet increased in association with elevated colonic Cld-2 expression (Figure 6B and 6C, Figure S5), maitake feeding normalized the expression of Cld-2 as well as purified vitamin D_2_-feeding. These results strongly suggest that vitamin D_2_ in maitake exerts protective effects against colonic inflammation by improving gut barrier function, which may result from Cld-2 regulation through alternative vitamin D signaling. The lack of a significant difference in plasma Ca^2+^ concentration among the diet groups also suggests that the different colitis indices among the groups were probably not due to the calcemic effect of vitamin D, but rather due to other actions of vitamin D in the colon.

Even though the plasma concentration of 25(OH)D_2_ in maitake- or vitamin D_2_-fed mice was approximately half of the 25(OH)D_3_ concentration in control mouse plasma, the biological responses of the rescued mice, such as the induction of vitamin D-responsive genes, were almost comparable to those of control mice. This is probably due to the difference in binding affinity to vitamin D-binding protein (DBP) between 25(OH)D_2_ and D_3_. It has been reported that 85-90 % of circulating 25(OH)D is chaperoned by binding to DBP, and less than 1 % remains in its free form [42]. A previous study demonstrated that mice raised on either vitamin D_2_ or D_3_ alone (1000 IU/kg) showed different profiles of the circulating form, with approximately 2-fold higher levels of free 25(OH)D in vitamin D_2_-fed mice than in vitamin D_3_-fed mice, whereas the total 25(OH)D concentration was almost equivalent in both mice. Under these conditions, vitamin D_2_-fed mice showed more osteogenesis than vitamin D_3_-fed mice [43]. Although 25(OH)D is absorbed into the renal proximal tubule cells in a DBP-bound form via megalin/tubulin-mediated receptor endocytosis for CYP27B1-mediated conversion to 1,25(OH)_2_D [44], 25(OH)D is also taken up by megalin/tubulin-negative cells in the free form, which includes monocytes [45, 46]. These studies indicate that vitamin D_2_ exerts higher biological activity than vitamin D_3_ in some cell types and physiological conditions, such as the immune system, through the lower binding affinity of 25(OH)D to DBP. Thus, the pharmacokinetic differences between D_2_ and D_3_ may result in different biological activities *in vivo*. In addition, we previously reported that 25(OH)D_3_ exerts an antiproliferative effect in human prostate PZ-HPV-7 cell lines, in which the generated 1,25(OH)_2_D_3_ is negligible [47]. Therefore, 25(OH)D_2_ itself may exhibit biological activities similar to those of 25(OH)D_3_ under certain physiological conditions; however, further studies are required to confirm this.

Meanwhile, the potential biological effect of dietary fibers such as β-glucan should be considered in maitake-fed mice in the present study. Edible mushrooms provide a valuable source of carbohydrates, proteins, vitamins, antioxidants, and minerals, while being low in fat and calories, making them a nutritious and healthy dietary option [48]. Therefore, future studies should explore the potential synergistic effects of these components in maitake mushrooms for the prevention of colitis. Maitake contains 3.8% water-soluble polysaccharides, of which 13% is β-1, 3/1, 6-glucan [49]. The polysaccharides including β-glucans are recognized as the main bioactive components responsible for the various health-promoting attributes of maitake [50]. In particular, the physiological functions of mushroom-derived β-glucans for gut health have been highlighted, most of which are mediated by the modification of gut microbiota profiles [51–54]. Regarding the interaction between β-glucans and gut microbiota, we previously demonstrated that soluble oat fibers, which contain β-glucan, exerted a protective effect on 2,4,6-trinitrobenzenesulfonic acid (TNBS)-induced colitis. This effect was mediated through an altered gut microbiota composition, increased butyrate production, and a subsequent rise in the percentage of colonic Tregs, which was consistent with the overrepresentation of *Faecalibaculum rodentium* [55]. Altered gut microbiota profiles may affect the gut immune system through microbe-mediated metabolites, such as short-chain fatty acids, that exert immune regulatory effects [56]. Additionally, vitamin D also acts as a modulator of intestinal barrier and immune system [57]. Therefore, the interactions between vitamin D and dietary fibers in maintaining intestinal immune homeostasis should be further examined in the future, although the potential adverse effects of dietary fibers during the active colitis phase should also be carefully considered.

Several strategies to prevent gastrointestinal disorders using mushrooms have been reported [58]. Extracts containing polysaccharides from *Hericium erinaceus* (*H. erinaceus*), a traditional medical mushroom, have shown attenuative effect in mouse colitis models induced by TNBS and DSS, primarily through anti-inflammatory and anti-oxidant mechanisms [58]. Clinical trials have also revealed that an *H. erinaceus*-based nutraceutical compound enhances the effectiveness of oral 5-ASA in mild-to-moderate ulcerative colitis [59]. While *H. erinaceus* is widely recognized as a medicinal mushroom known for its anti-inflammatory and immune-modulating properties, the potential of other mushrooms in the prevention of gut inflammation is also gaining attention. The findings in the present study emphasize the potential of vitamin D-mediated nutritional strategies for protecting against gut inflammation, with particular focus on the unique benefits of edible mushrooms compared to traditional medicinal mushrooms like *H. erinaceus.*”

In summary, we demonstrated the beneficial effects of dietary maitake intake on gut health, which may be partially mediated by maitake-derived vitamin D_2_. Although further analyses are needed to elucidate the mechanisms underlying the attenuation of colitis by maitake-derived vitamin D_2_, these findings indicate that maitake is a good nutritional source of vitamin D that compensates for vitamin D_3_-deficiency and attenuates colitis progression in the D_3_-deficient model. From the aspect of gut health, maitake can be much better than animal vitamin D-rich food, as synergistic effects with dietary fibers may improve gut conditions through the microbiota. Further evaluation of the health benefits of maitake may suggest novel approaches to prevent disease onset in the intestine and colon.

## Acknowledgement

We are indebted to HOKUTO Corporation for kind supply of maitake mushrooms. We would like to thank Editage for English language editing. This work was supported by a research grant from Hokuto Bioscience Promotion Foundation.

## Author Contributions

Responsible for conceptualization, data analysis, writing of the original draft: M. N. and S. I. Conceptualization and methodology: M. N. Data acquisition and data analysis: M. N., R. M., Y. K., S. C., Y. F. Writing original draft: M. N., S. I. Review & editing of the original draft: R. M., Y. K., S. C. Review and editing of the original draft, supervision : Y. F. Funding acquisition: M. N. All authors read and approved the manuscript for publication.

## Abbreviations

25(OH)D: 25-hydroxyvitamin D
1,25(OH)2D: 1α,25-dihydroxyvitamin D
CD: control diet
Cld-2: claudin-2
CYP: Cytochrome P450
DAI: disease activity index
DBP: vitamin D binding protein
DD: vitamin D_3_-deficient diet
DMEQ-TAD: 4-[2-(6,7-dimethoxy-4-methyl-3-oxo-3,4-dihydroquinoxalyl)ethyl]-1,2,4-triazoline-3,5-dione
DSS: dextran sulfate sodium
G. frondosa: Grifola frondosa
H&E: haematoxylin and eosin stain
HD: high dose of vitamin D_2_-fortified DD
HM: high dose of maitake-fortified DD
IBD: inflammatory bowel diseases
LD: low dose of vitamin D_2_-fortified DD
LM: low dose of Maitake-fortified DD
PTH: parathyroid hormone
VDR: vitamin D receptor

