## Supplementary material for "Attenuative effect of *Grifola frondosa* (maitake mushroom) on severe DSS-induced colitis in vitamin D-deficient mice": SI: Nishikawa et al_SI_241109_Rv1.docx

**Method S1. Gut permeability assay in colitis mice**

Mice fed the test diet for 6 weeks were subjected to DSS-challenge for 5 days as described in method section. They were fasted for 4h before the assay. Fluorescein Isothiocyanate (FITC)-dextran (4 kDa, Chondrex, Woodinville, WA, U.S.) was intragastrically administrated to the mice at a dose of 5.0 mg/kg. After 2h of the administration, the blood was collected from caudal vena cava under the isoflurane anesthesia and then the mice were scarified. After centrifuged of the blood at 2,000 × *g* for 10 min, the resultant plasma was diluted with twice volume of PBS. Fluorescent intensity of the samples was measured spectrophotometrically using Spark multimode microplate reader (TECAN, Männedorf, Switzerland) in black 96-well plates with wavelength at 490 nm and 530 nm for excitation and emission, respectively.

**Table S1. Composition of vitamin D-related ingredients of test diet used in this study**

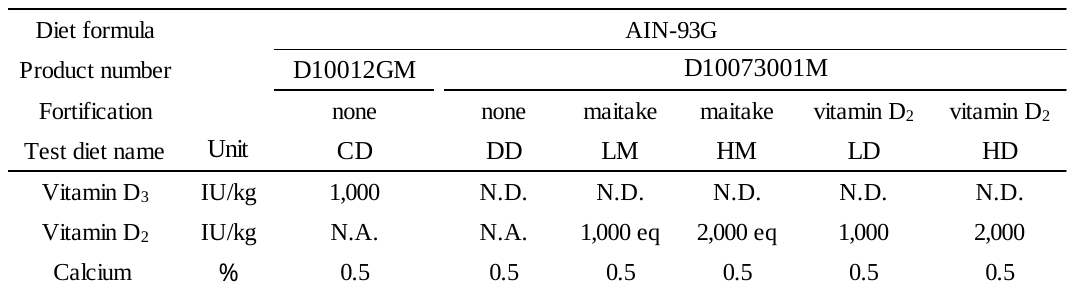

N.D.; not detected.

N.A.; not analyzed.

eq; equivalent concentration

**Table S2. LC-MS parameters for quantitative analysis of 25(OH)D_2_ and D_3_**

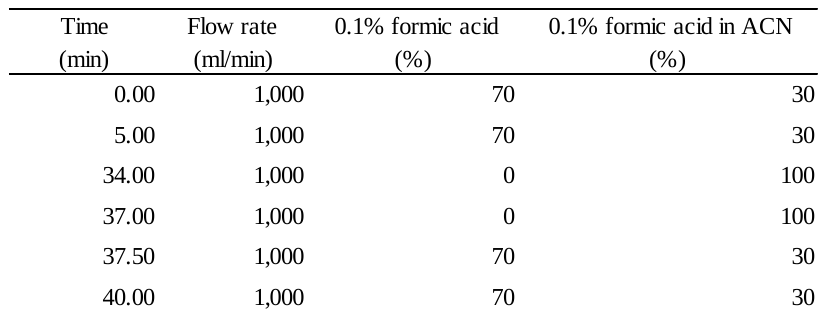
HPLC gradient program

Column; CAPCELL PAK CAPCELL PAK C18 UG120, 5 μm; 4.6 I.D.×250 mm (SHISEIDO, Tokyo, Japan)

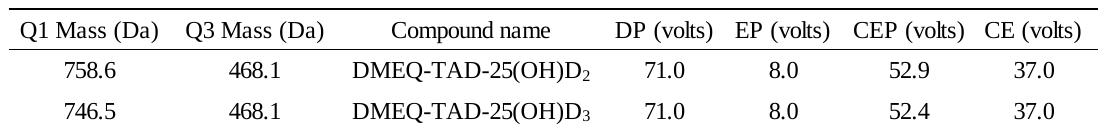
MS parameters

APCI-positive

**Table S3. Primer sequences for RT-qPCR**

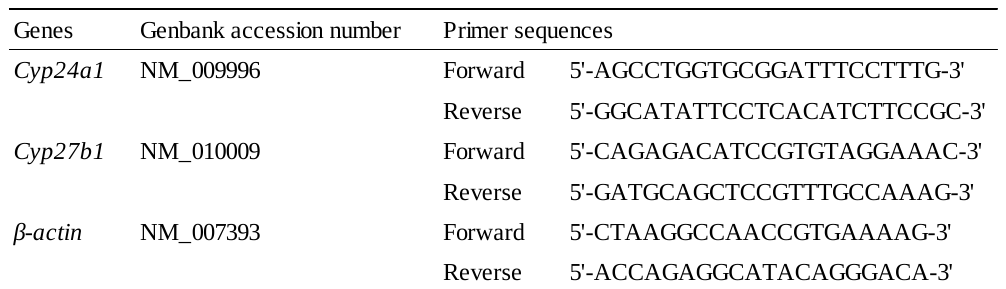

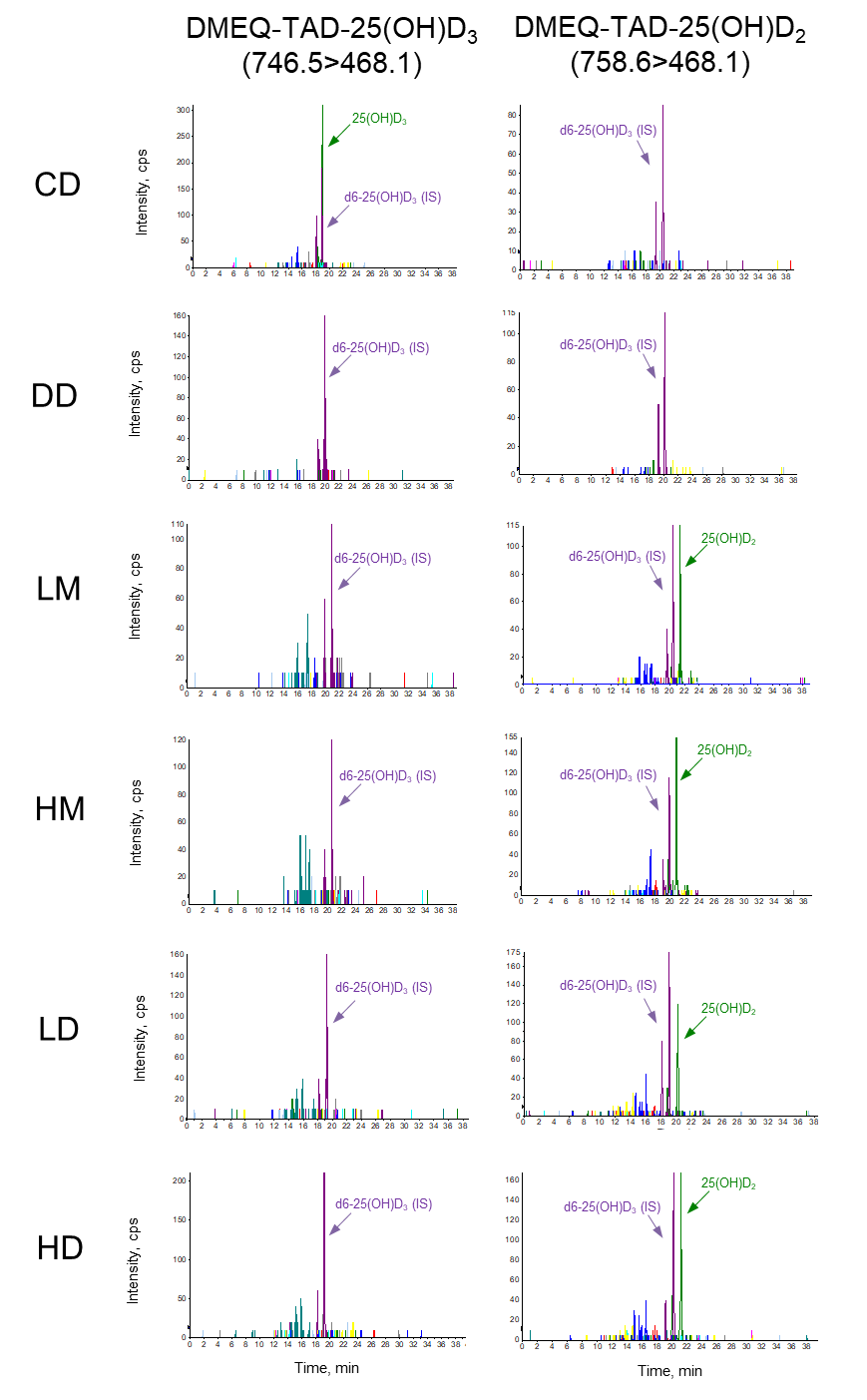

**Figure S1. TIC of the MRM chromatogram for 25(OH)D_2_ and 25(OH)D_3_**

| Duration (weeks) | Plasma 25(OH)D3 (ng/mL) |
| --- | --- |
| 0 | 26.68 |
| 1 | 1.55758 |
| 2 | 0.8 |
| 4 | 0.56 |

**Figure S2. Time-dependent effect of vitamin D_3_-deficient diet (DD) on plasma 25(OH)D_3_ concentration**

Mice was fed DD from 4 weeks of age. X-axis values indicate duration of DD‑feeding. The values are shown as the mean ±SEM (n=3 mice).

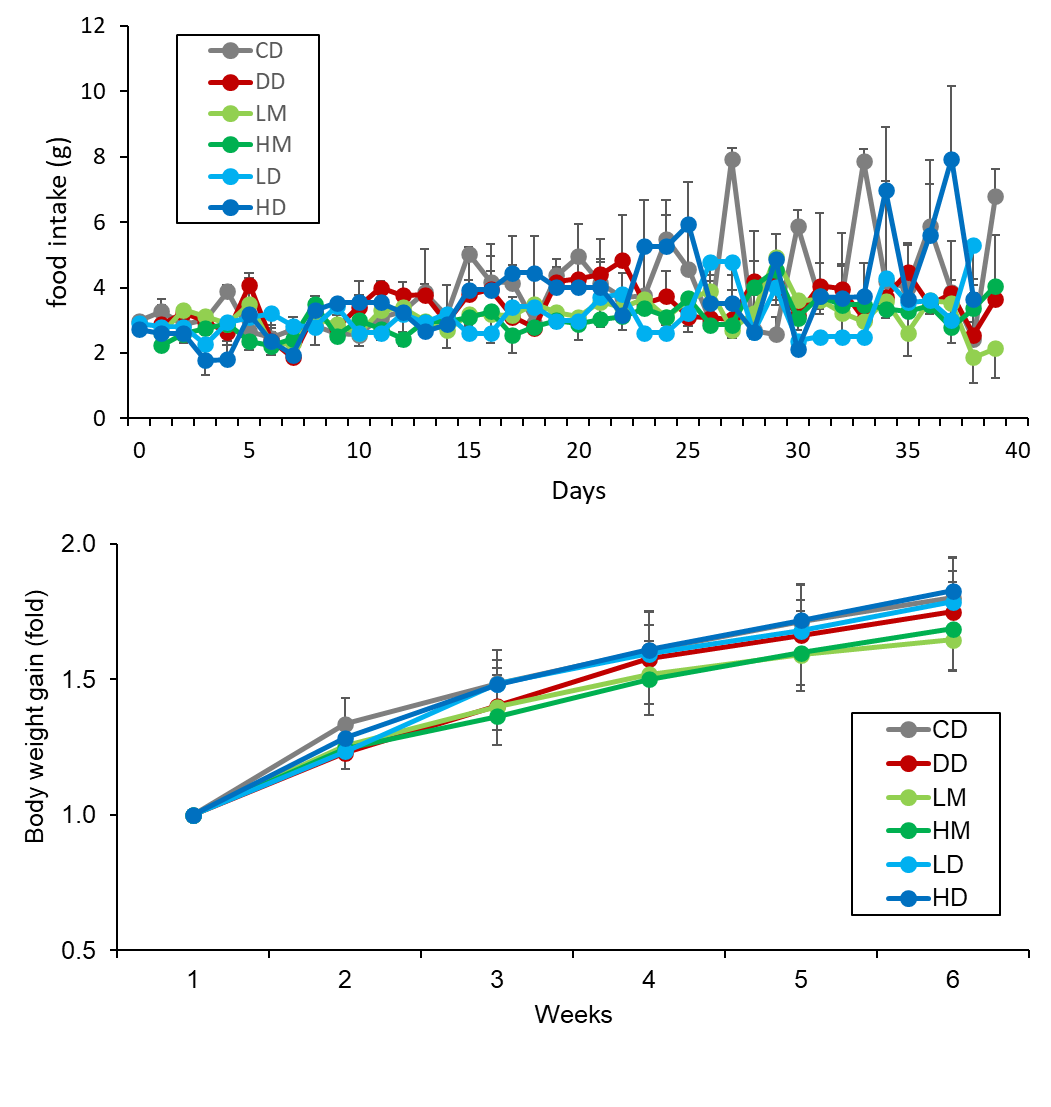

**Figure S3. Effect of vitamin D-controlled diet on food intake and growth during feeding period before DSS-challenge**

CD; control diet, DD; vitamin D_3_-deficient diet, LM; low maitake diet, HM; high maitake diet, LD; low vitamin D_2_ diet, HD; high vitamin D_2_ diet. The values are shown as the mean ±SEM (n=6-8 mice/group).

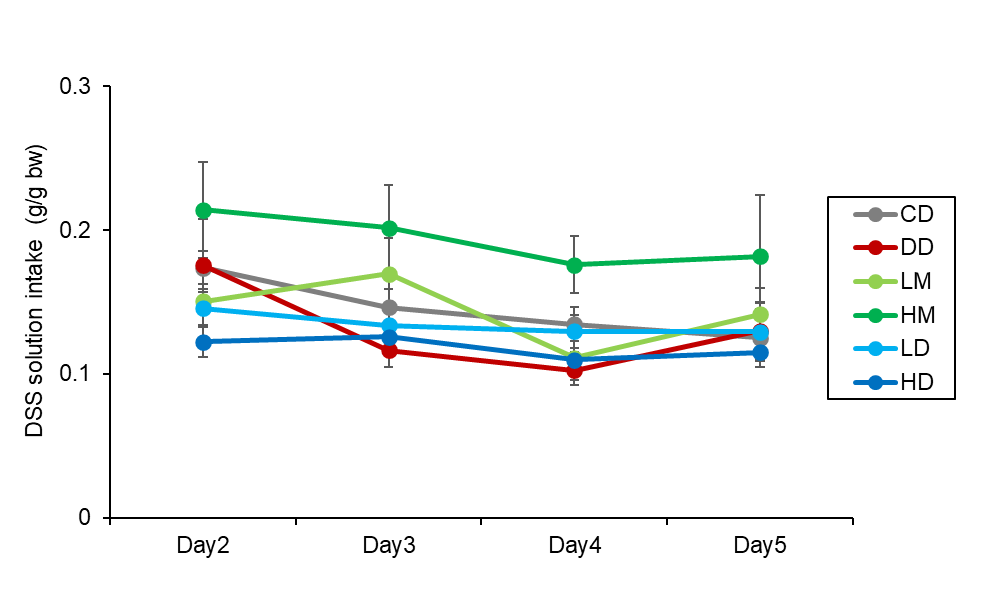

**Figure S4. Intake of DSS solution during DSS-challenge**

The amount of DSS solution intake is presented as daily decreased weight of the solution per bodyweight (g/g bw). CD; control diet, DD; vitamin-D_3_-deficient diet, LM; low maitake diet, HM; high maitake diet, LD; low vitamin D_2_ diet, HD; high vitamin D_2_ diet. The values are shown as the mean ±SEM (n=5-6 mice/group).

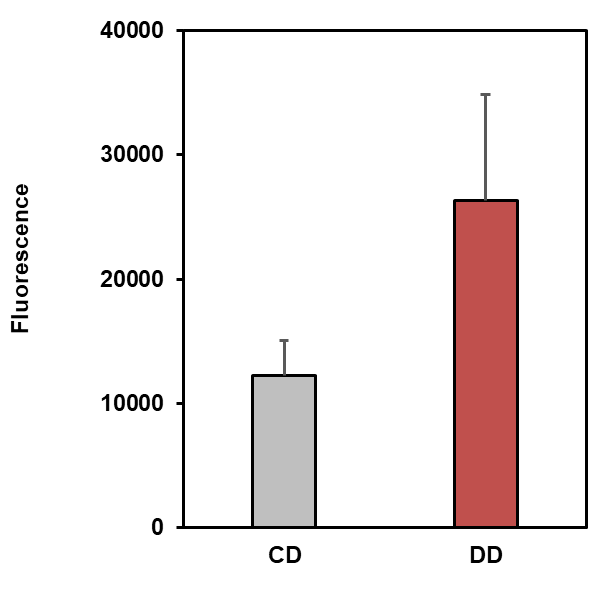

**Figure S5. Plasma concentration of FITC-dextran after 2h of intragastric administration.**

CD; control diet, DD; vitamin-D_3_-deficient diet. The values are shown as the mean ±SEM (n=3 mice/group).
